# *iMAX* FRET (Information Maximized FRET) for multipoint single-molecule structural analysis

**DOI:** 10.1101/2023.09.27.559821

**Authors:** Bhagyashree S. Joshi, Carlos de Lannoy, Mark Howarth, Sung Hyun Kim, Chirlmin Joo

## Abstract

Understanding the structure of biomolecules is vital for deciphering their characteristics and roles in biological systems. While current structural analysis techniques like nuclear magnetic resonance and X-ray crystallography excel in many aspects, they fall short in capturing comprehensive single-molecule information. To address this limitation and to better capture the heterogeneity and dynamic range of biomolecular reactions, there is a need for single-molecule structural analysis tools. To achieve this, we introduce iMAX FRET, a one-pot FRET-based single-molecule method integrated with geometrical 3D reconstruction, offering comprehensive *ab initio* structural analysis. Through the stochastic exchange of fluorescent weak binders, iMAX FRET allows simultaneous assessment of multiple spatial coordinates on a biomolecule within a few minutes of time to generate distinct FRET fingerprints for 3D structural profiling. We demonstrate a mathematical approach for *de novo* structural prediction using iMAX data, opening avenues for native biomolecule analysis. Furthermore, this method facilitates the investigation of conformational changes in individual molecules, illuminating single-molecule structural dynamics. Our technique has the potential to emerge as a powerful approach to advance our understanding of biomolecular structures.

## Introduction

Three-dimensional structure dictates the characteristics and functions of biomolecules^1^, and thus their analysis is fundamental to understanding their biological functions. Seemingly small perturbations—a single amino acid substitution, a change in local temperature, or an interaction with other molecules—can lead to a change in structure, which may ultimately lead to diseased cellular states^2-6^. As structure may differ from one protein molecule to another, analyzing the structures of individual single molecules and complexes is a prerequisite to understanding all cellular functions. However, traditional analysis techniques such as nuclear magnetic resonance and X-ray crystallography determine only the ensemble-averaged structure^7,8^ and thus are unable to capture the structure variation of individual molecules that may underpin crucial biological processes. Furthermore, they often impose artificial physical conditions during measurements (such as crystallization)^9-11 10^ and require complex methodology^12,13^. Single-molecule fluorescence resonance energy transfer (smFRET) and single-particle cryo-electron microscopy have emerged as cutting-edge techniques for interrogating structures of individual molecules. While complex workflow and heavy reliance on specialized experts of single-particle cryoEM hampers its cross-domain adaptability, smFRET is arguably less complex in its execution. smFRET can sensitively measure distances between fluorescent dye pairs attached to a biomolecule of interest in 2-10 nm range at sub-nanometer resolution and has been successfully used for conformational and kinetics analyses of biomolecule structures^14,15^. However, due to the complexity of signals, only one or two dye-labeled points in a single molecule can be tracked at a time^16^, precluding a comprehensive understanding of the three-dimensional perspective on the structure without prior knowledge of the molecular structure obtained by other means.

Extensions of conventional smFRET, which allowed observation of multiple distances between a single reference point and several other positions in a molecule of interest using a scheme of DNA exchange, were developed by several groups^17-19^. Although this multi-point analysis mitigated the limitation of the conventional smFRET, the necessity of a single reference point still implied that positions in three-dimensional space could not be triangulated for *de novo* structural reconstruction. We now present information MAXimized FRET (iMAX FRET), a one-pot experimental method to measure all possible mutual distance information between multiple selected points in a single molecule. Unlike hitherto reported FRET-based structural biology methods which require prior structural knowledge, iMAX FRET is the first method that enables *ab initio* structural analysis solely from smFRET data. Using our newly developed software pipeline, we show that iMAX FRET data can be used to determine up to six distances from four positions in 3D space, from which the conformation of a molecule can be reconstructed by geometrical modeling.

## Results

### The principle of iMAX FRET

*iMAX FRET* uses weak binders to rapidly assess multiple points in native biomolecules and heteromeric complexes (Fig. 1). In this work, we used short single-stranded DNA (ssDNA) as weak binders to exploit its programmable binding kinetics. Multiple positions of interest in protein, nucleic acid nanostructure, or multimeric complex were labeled with ssDNA molecules (hereafter called docking strand). Docking strands transiently hybridize with imagers – complementary DNA oligos in solution, labeled with either a donor or acceptor fluorophore (Fig. 1a). As these imager binding events occur stochastically and as each docking strand can serve as both the donor and acceptor binding site, all distances between the target positions can eventually be deduced from all the single-pair FRET events in which only a single donor and acceptor imager pair is bound to the target biomolecule. The lengths and concentrations of the imagers were tuned such that only one FRET pair is observed for a significant fraction of recording time. Collected FRET values are subsequently translated to distances, which are then fit together in a three-dimensional construct (Fig. 1b); all possible three-dimensional constructs using these lengths are generated, and the construct that violates the originally measured lengths the least is considered the correct fit. This approach obtains a per-molecule three-dimensional reconstruction without any prior knowledge of the identity or structure of the molecule; only basic geometry rules are applied.

**Fig. 1:**
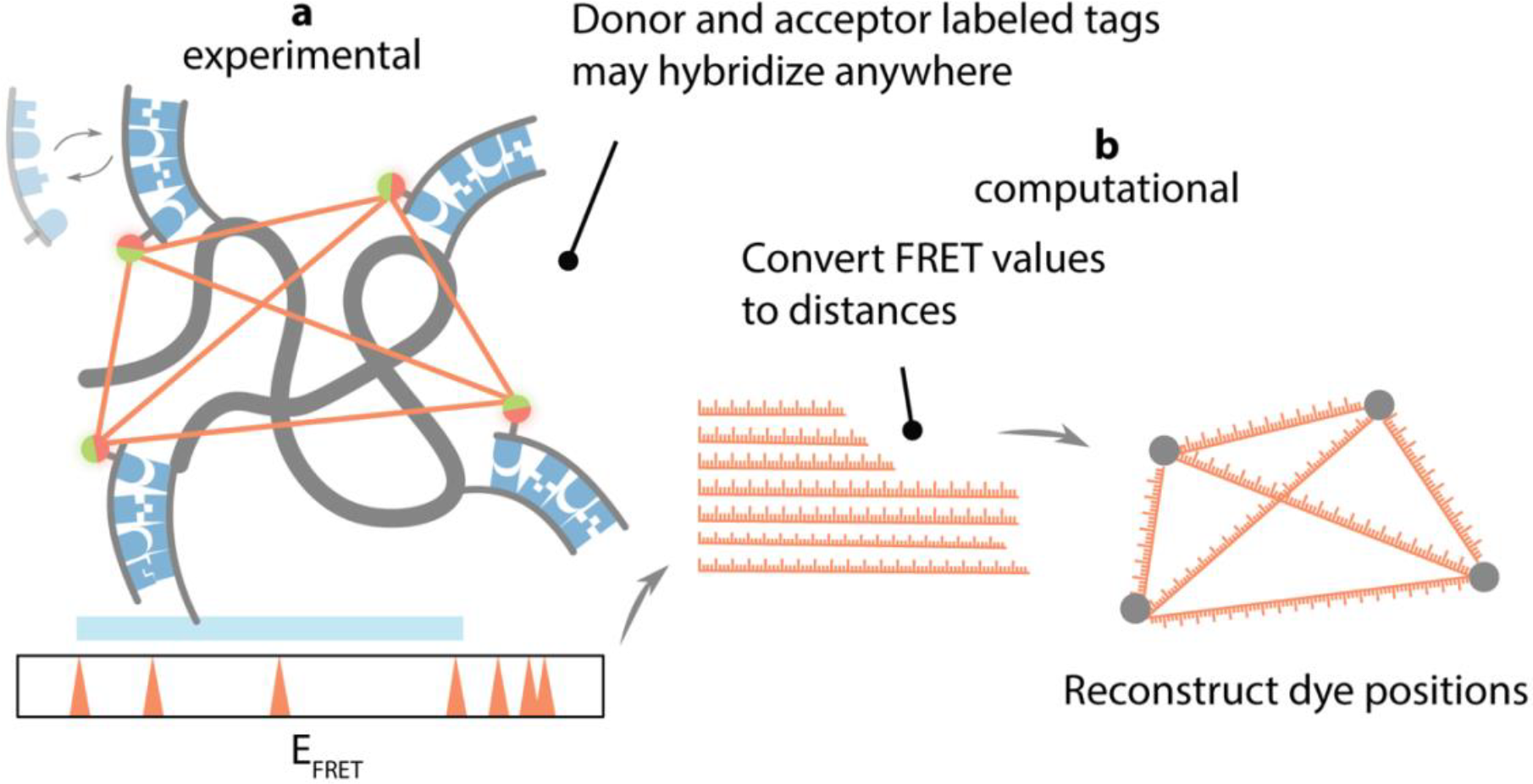
The general concept of iMAX FRET. **a**, <u>Experimental module:</u> A biomolecule consists of 2-4 coordinates carrying weak binder targets, here DNA docking strands to which cognate imagers can reversibly bind. The imagers are labeled with either donor or acceptor (green as donor and red as acceptor), and they both compete for the binding sites. Each successful FRET event has a particular FRET efficiency (E_FRET_) between two coordinates and, over time, all possible FRET efficiencies accumulate to give rise to the FRET histogram. **b**, <u>Computational module</u> -the apparent FRET efficiencies (E_FRET_) for single molecules are converted into distances. They are run through geometrical reconstruction to predict the most optimal fit for the structure. This designates the predicted structure calculated based on the apparent FRET efficiencies.

An advantage of iMAX FRET is its relative ease of implementation. A single round of standard two-color FRET measurement is sufficient to obtain all the structural information, whereas other single-molecule methods for multiple distance observation require the inclusion and observation of more dye colors, repeated measurements with probe exchange, or multiple sample preparations with different labeling schemes^16,19-23^. Furthermore, no orthogonal labeling schemes for different positions are necessary, as all positions can be labeled in the same way – with the same type of docking DNA strands, which are all stochastically visited by the same type of probes – here, imager DNA strands.

### iMAX FRET can delineate single-stranded DNA profile

As a first step, with a simple example of a ssDNA carrying multiple docking sequences, we sought to check the feasibility of the simultaneous multi-distance measurement with the one-pot stochastic probe exchange scheme. We prepared four ssDNA targets each of which contains two or three interspaced copies of an otherwise identical docking sequence at different positions, designated A, B, and C (Extended Data Fig. 1a, for sequences refer to Supplementary Table 1). Simultaneous binding to positions A and B – spaced 12 nt apart – was expected to give high FRET, positions B and C were spaced 19nt apart which should generate an intermediate FRET, and lastly, the summed 31nt distance between positions A and C should result in a low FRET signal (Extended Data Fig. 1a).

After immobilizing the ssDNA targets on a quartz slide glass via a biotin-streptavidin linkage, a mixture of donor and acceptor imagers of 8nt was added to the sample chamber. The 8nt imager, of which binding dwell time was τ_binding_=1.0 ± 0.1 s, was adapted from our previous study^19^. We added 10-fold excess of acceptor-labeled imagers over donor-labeled imagers to increase the probability of both fluorophores being present simultaneously. We then collected all the binding events from the time traces of individual molecules (Extended Fig. 1b) and built a histogram of the averaged FRET value per event (Extended Fig. 1c). All four DNA samples showed the expected FRET efficiencies of 0.73 ± 0.01, 0.52 ± 0.01, and 0.21 ± 0.02 for positions A, B, and C, respectively (Extended Data Fig. 1c). Notably, the DNA sample carrying all three docking sites showed all three peaks, confirming that our stochastic exchange scheme was capable of simultaneous multi-point assessment.

We noted that most binding events, however, showed FRET efficiency of 0.0 (star, peak area of ∼69 %) indicating that donor-only binding events are still dominant in the current experimental condition (Extended Data Fig. 1c). We anticipated that the acquisition of sufficient FRET events, i.e. simultaneous binding of a donor and an acceptor imager, required fine-tuning of the binding kinetics of the two imagers; event duration controlled by imager strand lengths, and event frequency controlled by concentrations of the two imagers. Using Monte Carlo simulations of our experiment (see Supplementary Methods) under different levels of concentration and binding dwell time of imagers, we inferred that 10-fold acceptor excess combined with longer acceptor binding times produced the optimal number of single FRET-pair events (Extended Data Fig. 1d and e). We extended the imager docking strands to 9nt and found that viable FRET events significantly increased compared to the same-length imagers (Fig. 2a-c, Extended Fig. 1f-g). This demonstrated that careful rational design of imager lengths, and hence dwell times, is pivotal in resolving multiple targets in iMAX FRET.

**Fig. 2:**
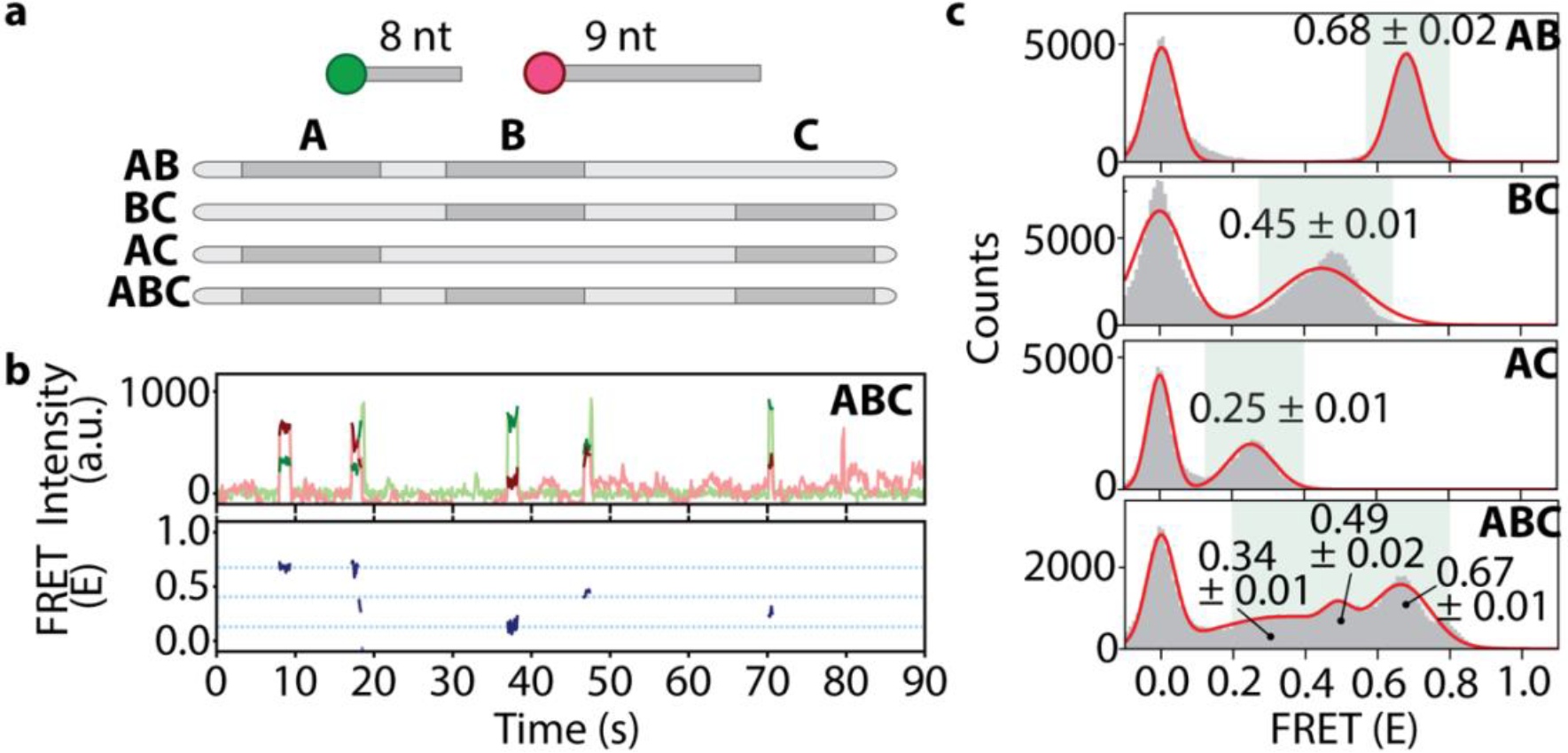
Resolution of three targets in linear DNA using iMAX FRET. **a**, Schematic representations of the linear DNA constructs. A, B, and C are the positions of identical docking sequences to which an 8nt donor- and a 9nt acceptor-labeled imager can bind. The molar ratio of the donor and acceptor strands were 1:10 (donor: acceptor). AB, BC, and AC are control constructs lacking either one of the three docking sequences, whereas ABC contains all three. The distances between A-B, B-C, and A-C are 12nt, 19nt, and 28nt, respectively. **b**, Representative single-molecule intensity vs. time trace (top panel) for the ABC construct (green for donor and red for acceptor intensities). Note that there are three different intensity peaks for the red i.e. acceptor intensity showing FRET events corresponding to successful donor-acceptor imager pair binding to A-C, B-C, and A-C docking sequences. The bottom panel shows the marked FRET efficiencies in blue. The highest blue line corresponds to A-B FRET, the middle line to B-C FRET while the lowest designates the A-C FRET event. **c**, Single-pair FRET event histograms from all molecules in a single field of view (grey bars). The mean FRET ± SEM is given for each peak in the histogram except the peak at 0.0 which corresponds to the donor-only binding events. Red solid lines are multi-Gaussian fit to the histograms. The FRET efficiency of each peak represents the distance between the designated docking sequences. Note the three peaks in the ABC construct corresponding to the three distances for A-C (0.34 ± 0.01), B-C (0.49 ± 0.01), and A-B (0.67 ± 0.01).

### iMAX FRET can resolve DNA nanostructures

To demonstrate iMAX FRET’s capability of *ab initio* three-dimensional structure determination, we analyzed a quadrangular DNA nanostructure outfitted with a docking strand at each angle (Fig. 3a, left). The six different distances in this nanostructure, referred to as D1 to D6 (Fig. 3a, right), could be probed with four identical docking strands in iMAX FRET. In this demonstration, however, we prepared each docking strand with a unique sequence for control purposes.

**Fig. 3:**
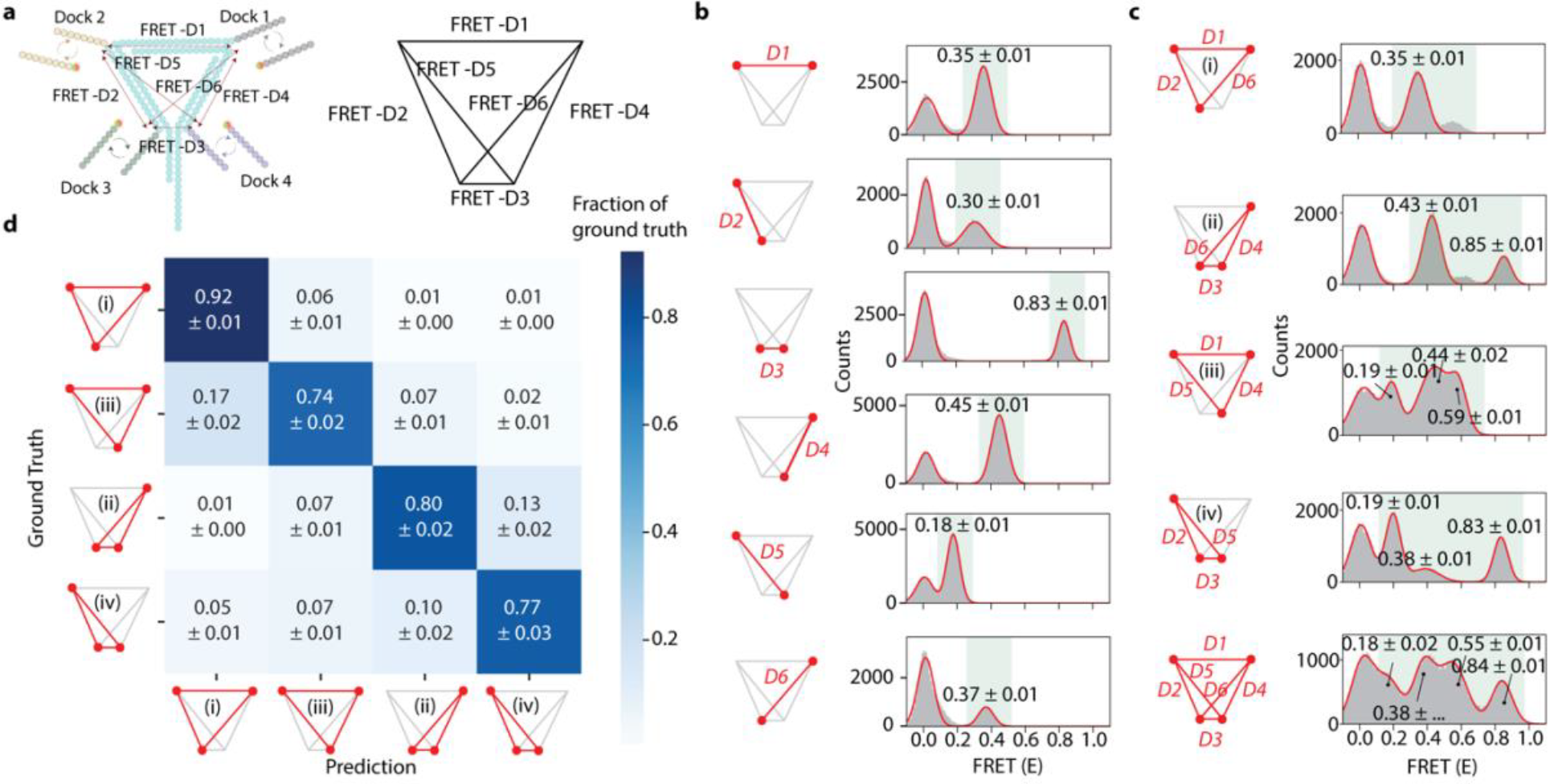
iMAX FRET provides structural analysis of a complex DNA nanostructure. **a**, Schematic representation of DNA nanostructure containing 4 overhangs of different DNA sequences which act as docking segments (Docks) for the imagers. As cognate imagers labeled with a donor or acceptor dye bind transiently to the Docks, FRET events occur in proportion to the distances between the Docks. 15 bp DNA length separates each pair of Docks 1-2 and 2-3, whereas the 13bp segment separates the Docks 1-4 giving rise to FRET distances of FRET-D1, D2, and D4, respectively. As a result, Docks 3 and 4 are situated very close to each other giving rise to FRET-D3. The Docks 2-4 and 1-3 also make a pair culminating in FRET-D5 and -D6, respectively. The right panel is the line representation of all the distances generated from the DNA nanostructure and will be used henceforth as a model figure. **b**, iMAX FRET histograms of each FRET distance D1 to D6, separately. The red lines signify the FRET distance, red dots represent the Docks. Note that D1, D2, D4, and D6 are similar while FRET D2 and D3 mark the extremes in either direction. The shorter length of DNA (FRET D4, 13bp) is reflected in slightly higher FRET efficiency (0.45 ± 0.01) as opposed to 2bp longer FRET D1 and D2 (0.35±0.01 and 0.30±0.01), respectively. This hints at the distorted nanostructure due to differential side lengths. **c**, iMAX FRET histograms for combinations of all 3 spatial points forming triangles (i-iv). The red lines signify the FRET distances, red dots represent the Docks. Note that triangle (i) has one mid-FRET degenerate peak due to the three overlapping distances of D1, D2, and D6. Triangle (iv) has one mid-FRET degenerate peak from D4 and D6, and a high-FRET peak arising from D3. The bottom structure contains four peaks with 2 degenerate peaks and 2 single peaks as a result of all the FRET distances D1-D6. **d**, Confusion matrix showing classification accuracy and error modes of a tree-based machine learning classifier trained to identify the four triangles (i-iv) on a single molecule level and tested on held-out molecules. Each row denotes which fraction of total molecules for a given ground truth class are ascribed to which class, where the diagonal denotes correct classifications (i.e. the per-class accuracies).

First, we probed each distance by adding two different imager strands simultaneously, which resulted in a single FRET peak per experiment (Fig. 3b). We found that D3 and D5 were well-discernible from the other distances (FRET efficiency mean ± standard deviation of 0.83 ± 0.01 and 0.18 ± 0.01, respectively). D1, D2, D4, and D6 generated highly similar FRET values (0.35 ± 0.01 and 0.30 ± 0.01, 0.45 ± 0.01 and 0.37 ± 0.01, respectively). Then, we increased the complexity by adding 3 imager strands, which allowed simultaneous analysis of three distances. Indeed, for each of the four possible triangles in this quadrangle, we could identify the expected number of FRET peaks (Fig 3c, panels i-iv). Triangle i (constructed from D1, D2, and D6) showed one major peak whereas for ii (D3, D4, and D6) two peaks overlapped, which was expected based on single-distance analysis results. Triangles (iii) and (iv), as expected, showed three peaks for (D1, D4, D6), and (D2, D3, D5), respectively.

To reconstruct the relative coordinates of dyes from per-molecule FRET values, each FRET efficiency *E* must first be translated to a distance *R*, following the sixth-power relation between *R* and *E*, and this requires estimating the Förster distance (*R*_*0*_) for our measurement conditions. This parameter combines the influence of dye and medium properties, and relative dye orientations. To estimate R0, we sought to utilize the known structure of double-stranded DNA. First, we prepared DNA nanostructures with one of the docking sites positioned at different locations along one arm of the nanostructure (Extended Fig. 4a-b) and probed two docks simultaneously. As expected, FRET values gradually decreased as we placed the variable docking site further from the fixed docking site (Extended Fig. 4c-e). We then probed an additional fixed docking site (i.e. three docks per nanostructure) and measured FRET distances for triangles, thus obtaining either two or three expected FRET *E* populations per molecule depending on the degeneracy in the FRET spectrum (Extended Fig. 4f). We could then estimate *R*_*0*_ by adjusting the value such that all triangles could be aligned by their fixed docks and with their variable dock in the position predicted from the dsDNA geometry (Extended fig. 4g and Supplementary Table 2; see also *DNA structure modeling for Förster radius fitting* in Supplementary Methods) Subsequently, FRET efficiencies were converted to distances using this fitted *R*_*0*_.

Having acquired the distances, the reconstruction of triangle coordinates is trivial as only one dissimilar triangle (i.e., ignoring rotation, translation, and reflection) can be constructed given the lengths of all three edges. Aligning and averaging triangle coordinates of all single molecules (Extended Fig. 2a), produced the shapes of the four triangle types (Extended Fig. 2b). To demonstrate that triangles reconstructed for single molecules contain sufficient information to allow recognition, we encoded the coordinates in a rotation-, translation- and reflection-invariant embedding^24^ and trained a tree-based machine learning algorithm to recognize each type. On held-out molecules, this classifier attained an accuracy of 74%, confirming that spatial reconstruction contains discriminative information (Fig. 3d). Most errors were made between triangles that were expected to show more similarity due to the nanostructure’s assumed symmetry (i+iii, ii+iv respectively). Even so, the classifier still correctly assigned classes to most molecules, thus we suspected that the nanostructure is not truly symmetric.

Finally, we probed all six distances simultaneously by adding four different imagers together (Fig. 3c, bottom plot). We observed four peaks. The highest peak at *E*=0.84 and the lowest at *E*=0.18 represented D3 and D5 respectively. The other two peaks were, however, not straightforward to assign due to the overlapping FRET values of the other four distances D1, 2, 4, and 6. Nevertheless, the broad peak at 0.38 could be assigned as a degenerate peak of D1, D2, and D6, while the peak at *E*=0.55 likely arose from D4. FRET values obtained from this experiment were then used to reconstruct a 3D quadrangle. In theory, 30 dissimilar quadrangles can be constructed given the lengths of all six edges. However, not all quadrangles can necessarily be built without violating the given lengths. We therefore wrote an analysis pipeline (see Supplementary Methods) that builds all possible dissimilar quadrangles and chooses the one for which the edge lengths are required to change the least to fit. We found that a 3D quadrangle could be constructed satisfying all distances without violating the FRET-derived lengths (Extended Fig. 3). Similar to the triangle measurements, this reconstruction indicated that the nanostructure has an asymmetric conformation, possibly reflecting the slight out-of-plane attachment positions of the docking strands due to the helical structure of the double-stranded DNA. Also, spatial hindrances due to electrostatic repulsion from the DNA nanostructure might have influenced the actual positions of the docking and imager strands, which could push the dyes further away from the attachment point depending on the local 3D geometry.

In summary, iMAX FRET could successfully demonstrate the structural analysis of up to 4 points in a complex DNA nanostructure, and we could predict and retrieve these structural identities with high accuracy based on FRET fingerprints and computational modeling. This demonstrates that we could expand the signal space to 6 peaks (considering degenerate peaks) in a one-pot reaction requiring less than 2 minutes without using solution exchanges^19^.

### iMAX FRET locates the biotin pockets in tetravalent and divalent streptavidin structures

As iMAX FRET is well-suited to determine the relative position of three or more points in space, we set out to study multimeric structures, which are difficult to analyze with traditional FRET due to the inability to control labeling with donor and acceptor fluorophores of subunits within a multimeric protein^25^. Structural analysis of multimeric proteins by other techniques, including mass spectrometry, often requires complex stabilization using chemical linkers or cross-linking^26-29^. In contrast, iMAX FRET can be applied on native complexes. Moreover, ligand-binding multimers present a unique possibility for iMAX FRET. For example, we can use docking strand-conjugated ligands to probe the positions of their binding pockets. We chose streptavidin as our model protein, as it contains four pockets for biotin. This also allowed us to indirectly immobilize streptavidin to a surface, by occupying one of its pockets with an immobilized biotinylated docking strand (Extended Fig. 5a). The other pockets were occupied by docking strands added in solution.

Streptavidin is a tetramer organized in a tetrahedral (D2) symmetry with four biotin-binding pockets (Extended Fig. 5b) First, to derive single distances from four binding pockets, we measured two divalent streptavidin mutants – 1,3 *trans* and 1,2 *cis* which have only two active biotin binding pockets^30^ (Fig. 4a and b). As expected, a high FRET peak (0.89 ± 0.04) was observed for 1,2 *cis* and a mid-FRET peak (0.56 ± 0.02) for 1,3 *trans* (Fig. 4a and b). Changing the dye positions from one end to another of the imagers proportionately reflected the changes in the FRET values, showing the ability of iMAX FRET to pinpoint the biotin binding pockets accurately (Extended Fig. 5c and d). Subsequent iMAX FRET analysis on the wild-type streptavidin with four active binding pockets showed three different FRET efficiencies of 0.28 ± 0.02, 0.58 ± 0.04, and 0.94 ± 0.04 seen for pockets 1, 2, and 3, respectively (Fig. 4c). Although, in general, six distances were expected from four points, the symmetric tetramer structure of streptavidin could exhibit only three peaks due to degeneracy. Nevertheless, by using these FRET values, we were able to reconstruct the relative positions of the binding pockets (Fig. 4d). The reconstructed 3D spatial coordinates fit the known streptavidin structure gratifyingly well, accounting for a realistic average linker length of 1.8nm, and showed limited variability over 1000 bootstrap iterations (standard deviation of 2.75Å averaged over all four positions, Fig. 4e). This confirms that iMAX FRET is capable of extracting three-dimensional features from multimeric proteins without the aid of complementary methods or additional information on the target.

**Fig. 4:**
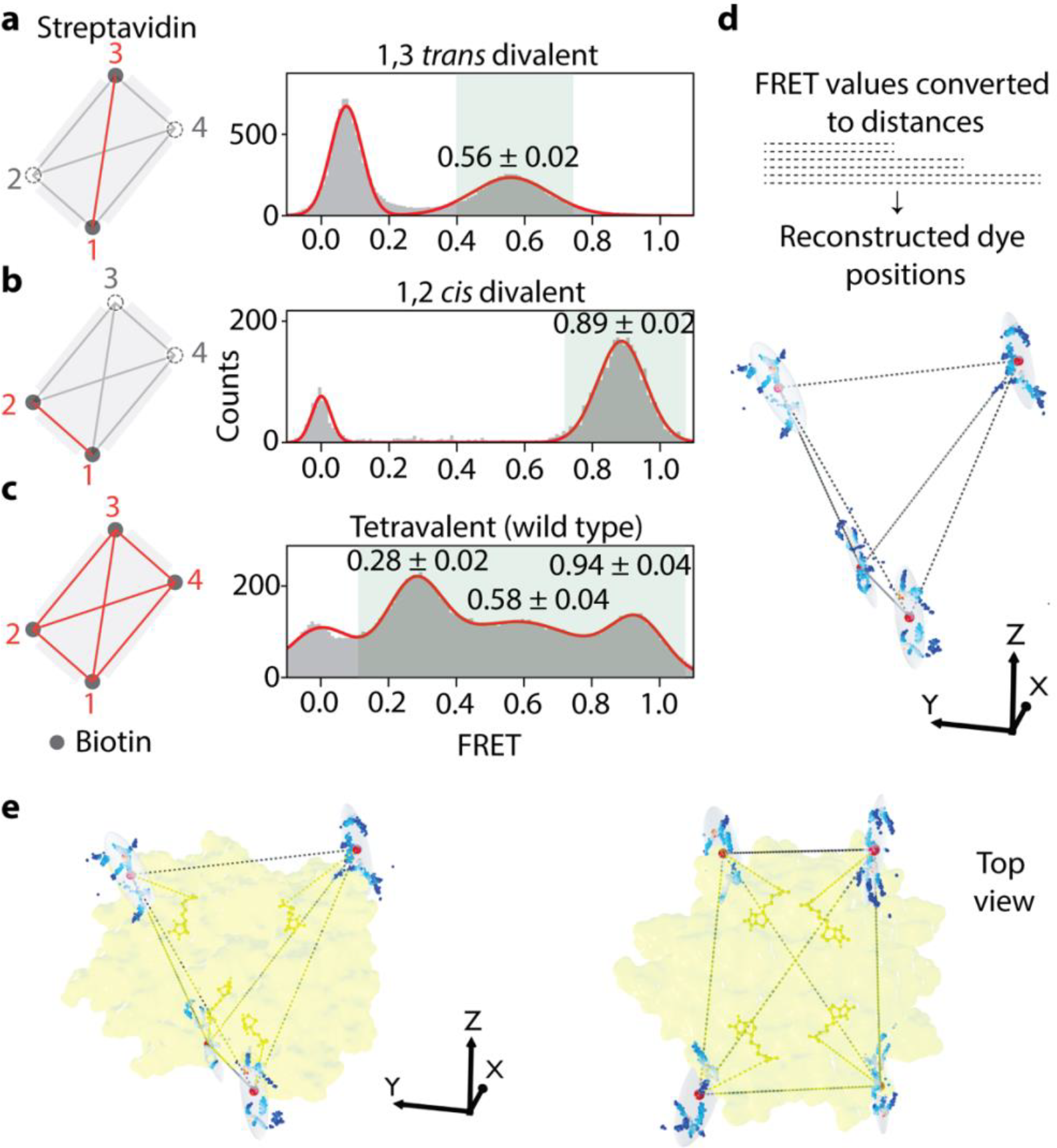
iMAX FRET-based structural analysis of streptavidin complexes. **a**, The mutant *1,3 trans* divalent streptavidin (PDB ID: 4BX6) that can bind biotin (grey dots) to only two binding pockets, whereas the other two are mutated to abrogate the biotin-binding (dashed circles). Upon binding of an imager, the dye is facing towards the binding pocket. The distance between the bound biotins (red line) shows the FRET efficiency of 0.56 ± 0.02 in the histogram. **b**, The mutant *1,2 cis* divalent streptavidin (PDB ID: 4BX5) that can bind biotin as shown. The distance between the bound biotins (red line) shows a high FRET efficiency of 0.89 ± 0.02. **c**, The wild-type tetravalent with four active biotin-binding pockets. Hence, it can give rise to six distance possibilities [n(n-1)/2]. However, streptavidin is a symmetrical molecule, hence shows three degenerate peaks, each peak corresponding to two overlapping peaks. **d**, The FRET values are converted to six distances, and a structure is reconstructed for four biotin-binding pockets. Average positions for 1000 bootstrap iterations over all molecules are shown as dots (colored by density), the mean position is shown as a large red sphere and ovals report one standard error intervals (on average, 2.75Å). **e**, The reconstructed structure is fitted into the reported (PDB ID: 2IZF) crystallographic streptavidin structure (yellow). Note that all four biotins can be fitted into the biotin-binding pockets with high accuracy.

### iMAX FRET has a potential for studying permanent protein conformational changes

Finally, we explore the possibility that the stochastic DNA probe-based detection scheme used in iMAX FRET is also compatible with studying conformational changes of proteins without disturbing their activity. Many proteins undergo profound conformational changes upon binding to a ligand. A well-known example is substrate binding domain (SBD)^31^ which captures extracellular substrates and delivers them to transporters. We focused on the SBDs of one such transporter, GlnPQ from *Lactococcus lactis*, which are involved in amino acid sensing and import of asparagine and glutamine^32,33^. Here, we attempted to detect the open-to-closed conformational switch after ligand binding to SBD2 protein^34,35^.

We prepared a wild-type protein that can bind glutamine and asparagine and a null mutant that does not bind any ligand as a control^35^ (Extended Data Fig. 6a). For DNA labeling, two cysteines were inserted into both proteins at strategic positions each located at one of the two lobes in SBD2 (Extended Data Fig. 6b). These modifications are known to have no adverse effects on their function^35^ and the distance between these two positions undergoes a significant change after a ligand binding according to the crystal structures^34^. Indeed, observed FRET increases from 0.25 to 0.40 and from 0.25 to 0.30 upon binding of Glutamine and Asparagine, respectively (Extended Fig. 6c). The smaller FRET shift with Asparagine reflected the fact that SBD2 undergoes a higher conformational change when bound to Glutamine compared to Asparagine^35^. In contrast, we did not observe a FRET shift from the mutant, confirming the FRET shift is indeed induced by ligand binding (Extended Data Fig. 6d). We conclude that iMAX FRET with the stochastic DNA probe exchange method can be applicable to dynamic structural analysis of proteins as a response to stimuli.

## Discussion

Here we presented iMAX FRET, a novel structural analysis tool to probe multiple pairwise distances by using high-resolution smFRET and weakly interacting probe scheme. By directly integrating it with geometrical modeling for structural prediction, iMAX FRET enables the assessment and prediction of molecular structures based on their FRET fingerprints with the ultimate sensitivity of a single molecule, thus opening avenues for new *ab initio* structural prediction and conformational dynamics studies.

iMAX FRET has many advantages over established techniques. *a) Short time required for measurement*: As we use the stochastic exchange scheme for probing all possible points in a molecule with otherwise identical probes, this one-pot method cuts down the imaging time considerably as compared to other DNA hybridization-based imaging techniques ^19,23,36^. The probe-labeled samples can be practically prepared in 24 hours ^19^ while weak-binder based fluorescence measurement takes as little as two minutes. Thus, the structural analyses of protein and complex mixtures can be obtained multiple times faster than other structural analysis techniques. *b) Ease of sample preparation*: iMAX FRET overcomes challenging sample preparation or crystallization as required for CryoEM and X-ray crystallography. As only picomolar-range quantities are required for the analyses, it is possible to analyze the most precious samples (e.g. patient materials). The stochastic nature of iMAX FRET measurement makes the sample preparation easier as only one type of attachment chemistry is used for all the docking sites. If necessary, orthogonal labeling and pull-down methods such as His-tag or N-terminus labeling^37^ may be used to make the technique more easily accessible and applicable.

iMAX FRET allows probing of multiple distances in a nano object, including complex DNA nanostructures, proteins, and heteromeric complexes of biomolecules. It paves the way for studying the static and dynamic structural analysis on challenging multimeric proteins such as transcription factors and transmembrane proteins. iMAX FRET has the potential to provide quantitative information on the species abundance of multimers and their characteristics in a complex mixture of homo and heteromers. Thus, it can replace cumbersome biochemical assays used to delineate the differential populations of homomers and heteromers present in a particular solution.

Even with its empirical simplicity, iMAX FRET can provide useful 3D structural information of biomolecules. Yet, certain problematic proteins, such as those 1) with too large or complicated structures, 2) with too low or high number of possible labeling points, or 3) with too much degeneracy in the measured distances, may be analyzed over multiple rounds of measurements by utilizing the programmable nature of a probe. Further developments should, therefore, include identifying and evaluating widely appliable methods for the orthogonal labeling of docking strands or the use of other weak binders that do not require protein labeling. A logical first step of labeling to this end is the targeting of cysteine residues^38^, which are rare amino acids and are primarily surface-exposed, making them immediately amenable for use in iMAX FRET. Our technique can also be extended to other amino acids such as lysine of which conjugation chemistry is well established^39^. This extension may open the possibility to structures that are difficult to assess using CryoEM and X-ray crystallography or predict with Alpha-fold due to their intrinsic disorder^40^ or propensity to irregularly aggregate, or change structure during complex experimental workflows.

## Author Contributions

B.S.J. and C.J. initiated and designed the project. B.S.J. and S.H.K. designed and performed the experiments. I.W. expressed and purified the proteins. S.H.K. designed and ran FRET efficiency extraction workflows and Monte Carlo simulations. C.L. designed and ran 3D modeling and structure classification pipelines. M.H. provided the streptavidin mutants. B.S.J, C.L., S.H.K., and C.J. wrote and edited the manuscript. All the authors read and approved the manuscript.

## Acknowledgments

We thank Mike Filius for his generous help with the DNA constructs and overall scientific advice. C.J. is supported by The Netherlands Organization for Scientific Research (NWO) (Vici), the European Research Council (an ERC Consolidator grant, 819299), Basic Science Research Program (NRF), and Frontier 10-10 (Ewha Womans University). M.H. acknowledges funding from the Biotechnology and Biological Sciences Research Council (BBSRC, BB/I006303/1).

## Data availability

The data supporting the findings of this study are available in the article and its supplementary information. Any additional data is available from the main corresponding author upon reasonable request.

## Code availability

The iMAX FRET analysis pipeline and additional custom analysis codes are documented and freely available at https://github.com/cvdelannoy/iMAX-FRET.

## Materials and Methods

### Protein expression and purification

Divalent streptavidins were expressed in *Escherichia coli*, refolded from inclusion bodies, and purified by ammonium sulfate precipitation and ion-exchange chromatography, as reported in the original paper^30^. Tetravalent recombinant (wild-type) streptavidin was procured from Thermo Scientific. Plasmids encoding SBD2 (T369C/S451C) and SBD2 (T369C/S451C/D417C)^34^ were a generous gift from Prof. Bert Poolman (Department of Biochemistry, University of Groningen, The Netherlands), and the proteins were expressed and purified using the reported protocol ^34^.

### Protein labeling

Cysteine labeling was carried out as reported previously^34^ with slight modifications as follows. Cysteine residues of purified proteins (25uM in the total volume of 50ul In PBS) were reduced with 50mM Tris(−2-carboethyl)phosphine (TCEP) at 40-fold molar excess for 30 minutes. Excess TCEP was removed with ZebaTM Spin desalting columns 7kDa MWCO (ThermoFisher) as it may interfere with the Maleimide reaction^41^. The proteins were then labeled with 25-fold molar excess monoreactive maleimide-Dibenzocyclooctyne (DBCO) (Sigma Aldrich) in Phosphate Buffered Saline (PBS) pH 7.4 overnight at room temperature. Excess maleimide-DBCO was removed with Zeba columns and reacted with 10-fold molar excess (ratio 1:10, cysteine to linker) of monoreactive Azidobenzoate-(5’) functionalized DNA in PBS pH 7.4 and incubated overnight at room temperature.

### Single-molecule Setup

All iMAX-FRET measurements were performed on a custom-modified prism-type TIRF microscopy setup built around an inverted fluorescence microscope (Nikon, Ti2e)^42^. For illumination of samples immobilized on a quartz slide surface, a 532 nm diode-pumped solid-state laser and 640 nm diode laser (Oxxius, L6Cc) were directed to the surface with an incidence angle below the critical angle via a prism installed above the slide. Fluorescence signals of Cy3 and Cy5 dyes collected by an objective lens (Nikon, CFI Plan Apochromat VC 60X WI) placed below the quarts sample chamber were spectrally divided by a dichroic mirror (Chroma, T635lpxr) after removing scattered laser light by a laser blocking filter (Semrock, NF03-405/488/532/635E-25). The fluorescence signals were further cleared by bandpass filters (Chroma, ET585/65m for Cy3 and ET655LP for Cy5) and imaged on a sCMOS camera (Photometrics, PrimeBSI). The two lasers were operated with a trigger signal generated by the sCMOS camera for ALEX illumination scheme^43^. All the instruments were controlled by using commercial software (NIS elements, Nikon).

### Single-molecule flow cell preparation and data acquisition

All single-molecule FRET experiments were performed at room temperature. The flow cells were prepared using our published protocol^19^. Briefly, quartz slides (G. Finkerbeiner Inc) were etched using acidic piranha and passivated with polyethylene glycol (PEG) to minimize any non-specific binding of molecules. mPEG-SVA and PEG-Biotin (Layson Bio) were used for the PEGylation. 50ul of 0.1mg/ml streptavidin (Thermofisher) was incubated into the flow channel for 5min. Excess was removed using 100 μL T50 (50mM Tris-HCl, pH 8.0, 50 mM NaCl). Next, 50 μL of 100pM biotinylated samples was introduced and incubated for 5min in the channel: linear DNA (Fig. 2), triangles (Fig. 3), or biotinylated Anti-His antibody (Extended Data Fig. 5c and d). Unbound molecules were washed away with 100 μL T50. 100 μL of 10 nM donor labeled imager strands and 100 nM of acceptor labeled imager strands against the sequences under investigation were injected in imaging buffer (50 mM TrisHCl, pH 8.0, 500 mM NaCl, PCD (Merck), PCA (Merck) and 1 mM 6-hydroxy-2,5,7,8-tetramethylchroman-2-carboxylic acid (Trolox) (Sigma). See Supplementary Table 1 for the full list of docking and imager strands.

Generally, for single-molecule studies, immobilization is carried out by biotin-streptavidin interactions^42^. However, it is highly difficult to precisely control the number of biotin molecules on the traditionally passivated surfaces (with Biotin-PEG). This raises the possibility of 2 or more binding pockets of streptavidin being occupied by biotins on the slide, leaving only one or two for actual fingerprinting. Thus, we modified the immobilization strategy for tetravalent and divalent streptavidin experiments (Fig. 4): Quartz slides were sonicated for >15min in Acetone, Methanol, and finally 1M KOH with washes with MilliQ in between. Next, the slides were flamed using a burner to remove organic residue if any, and immediately placed back in MilliQ. Finally, the slides were dried using a nitrogen blowgun and used for making the flow cell as explained above. The unused slides were stored at RT. 50ul of 1mg/ml of BSA-Azide (Click chemistry tools, 1535) was incubated with 15ul of 100uM (5’) DBCO-DNA-Biotin (3’) overnight in the dark at room temperature. 10nM of the resultant BSA-DNA-Biotin was added (50ul total volume) to the flow-cell and incubated for 10min. Excess BSA and free DNA were removed with 100ul T50. Next, 50ul of 1nM tetravalent or divalent Streptavidin was added to the channel and incubated for 5min. The excess was washed with 100ul of T50. Next, 100nM biotinylated docking strands were added to the flow cell and incubated for 30 min to ensure the labeling of all the streptavidin pockets. Unbound DNA was washed away with 100 μL T50. Following, 50 μL of 10 nM donor-labeled imager strands and 100 nM of acceptor-labeled imager strands prepared in the imaging buffer were injected into the flow cell.

### Single-molecule fluorescence and FRET data analysis

The data collection and analysis were performed in multiple steps as reported previously^19^. A custom Python script was used to extract time traces of individual molecules from a sCMOS image collected at 0.1s exposure time per frame. Two-state K-means clustering algorithm were applied to the Cy3 and Cy5 fluorescence intensity traces to detect individual binding events of fluorescence imager strands. In order to ensure accurate results, binding events lasting for three or more consecutive frames were selected for further analysis. FRET efficiencies were calculated for each imager strand binding event and used to construct the FRET kymograph and histogram. From the events in which the acceptor probe dissociated or photobleached before the donor probe, we calculated the beta (leakage) and gamma correction factors for accurate FRET efficiency calculation following the method reported in a previous study [Biophysical Journal 99, 961–970]. Gaussian mixture modeling was applied to automatically classify populations in the FRET histogram. The Python-based automated analysis code can be freely accessed at the following link: https://github.com/kahutia/transient_FRET_analyzer2.

