## Supplementary Information for "*iMAX* FRET (Information Maximized FRET) for multipoint single-molecule structural analysis"

### Extended Information

#### *iMAX* **FRET** (**I**nformation **M**axi**m**ized FRET) for multipoint single-molecule structural analysis

#### Supplementary Methods

##### Monte Carlo simulations

In iMAX FRET, which has multiple identical docking sites, the chance of having single-pair FRET events, i.e. simultaneous binding of one cy3- and one cy5 probes, largely depends on the probe binding kinetics. Experimentally, the binding frequency and binding dwell time of a probe can be controlled by the concentration and the length of the DNA probe, respectively. To find the optimal condition that maximizes the chance of having FRET events, we carried out series of Monte Carlo simulations at various kinetic rates. We defined a system with three docking sites each of which had three states of 1) probe unbound, 2) cy3 probe bound and 3) cy5 probe bound states. Given transition rates of the two probes, each docking site of the system was allowed to freely transit between states 1 and 2 or states 1 and 3, but not between 2 and 3. Each simulation ran for 1-million-time steps from which we typically observed >5000 transitions. We then selected events in which the system entered into the single-pair FRET emitting state, in which only one cy3 and cy5 probe were bound among the three docking sites. After removing events that lasted shorter than three consecutive time steps, the number of the selected single-pair FRET events and the total time spent of the system in them were studied to understand the effect of probe binding kinetics. The simulation code was written in Matlab and freely available upon request.

##### Structure prediction and classification

A computational pipeline for the reconstruction of 3D-shapes and shape classification was implemented in Python 3.9. Briefly, the number of dyes is determined from the number of FRET efficiency values, which are translated to distances. Distances are used to construct all distinct distance matrices (*D*) using pre-computed index matrices. Each distance matrix is then converted to a coordinate matrix as follows[1]. We construct the Gramm matrix (*M*),

$$M_{ij} = \frac{D_{1j}^2 + D_{i1}^2 - D_{ij}^2}{2}$$

where *i,j* are row and column index respectively. After eigenvalue decomposition,

$$M = USU^T$$

the coordinate matrix *X* can be calculated by sorting U and S by descending order of eigenvalue size, taking the first 3 columns of *U* (*U*[:, :3]) and first 3 eigen values (*S*[:3]) and calculating:

$$X = U[:, :3]\sqrt{S[:3]}$$

Poorly fitting distance matrices generate negative eigenvalues and are excluded. Finally, the remaining coordinate matrices are calculated back to distance matrices, and the coordinate matrix for which distances are closest to the original FRET efficiency-derived distances is returned. The algorithm was implemented in numpy (v1.21.5)[2] with distance matrix calculation as implemented in scipy (v1.8.0)[3].

Numerical embedding of 3D shapes for classification was done using the Geometricus package (v0.3.0)[4]. Embedded coordinates were concatenated to the FRET fingerprint, after which a boosted tree classifier implemented using the XGBoost package (v.1.6.1)[5] was trained and tested on the data using a 10-fold cross validation scheme. The analysis code is freely available at https://github.com/cvdelannoy/iMAX-FRET.

**Preparation of the custom DNA nanostructure for Förster radius fitting and classifier applicability analysis**

The position of docking site 2 was changed to three different locations (Extended Fig. 4a) using click chemistry. To achieve this, alkyne handles were introduced into the DNA backbone at three different locations one at a time in separate constructs. The docking strand for site 2 was designed to contain an azide handle at its one end. The alkyne and azide-containing DNAs were reacted using copper-click chemistry. The clicked DNA products (cyan box, Extended Fig. 4b) were gel purified and then the triangles were assembled to generate three structurally similar nanostructures (Extended Fig. 4a, bottom left). The positional changes between the (variable) docking site 2 and the fixed docking site 3 were reflected in the FRET values (Extended Fig. 4c and d), while it remained constant for all the triangles for the undeviating distance between docking sites three and four (Extended Fig. 4e). We could similarly recapitulate the change in FRET values in three coordinates i.e. docking sites two, three and four (Extended Fig. 4f).

**Conversion of FRET efficiency into the distance R**

The following sixth-power relation between *R* and *E* was used to calculate the distance based on the experimentally acquired FRET efficiency.

$$E = \frac{1}{1 + (R/R_0)^6}$$

The Förster radius ($R_0$), a parameter that combines the influence of dye and medium properties, and relative dye orientations, was fitted using the above custom DNA construct with dyes positioned at known locations along one DNA arm (Extended Fig 4g).

**DNA structure modeling for Förster radius fitting**

The Förster radius ($R_0$) denotes the dye distance at which the FRET efficiency is 0.5 and constitutes an essential parameter for the accurate calculation of distances from FRET efficiencies[6]. It factors in dye quantum yields and relative orientations, and the refractive index of the medium. In many applications, it suffices to approximate this value as a constant, however in structural biology, this may lead to unacceptable discrepancies with actual distances, as the effect of local environment and setup is ignored. Here we have used an elegant experiment to determine $R_0$, using our DNA nanostructure. Briefly, we measure FRET efficiencies for four triangles, created by click-chemistry (detailed in Extended Fig. 4).

To determine the Förster radius for our experiments, a single side of the DNA nanostructure was outfitted with clicked docking strands at positions 4, 7, and 15$^{th}$ base from a reference position. FRET efficiencies between clicked docking strands, the reference position, and a third position at one of the other angles of the nanostructure were then measured. We then used a parameter optimization approach with a tree-based Parzen estimator (TPE) implemented in the hyperopt package (v.0.2.7)[7] to estimate the Förster radius. Briefly, this algorithm generates randomized proposals for all one or more variable parameters within given ranges and chooses the combination that minimizes the objective function. The TPE constrains the parameter space based on objective values of previous rounds so that the next guess is more likely to return a lower objective value. Using this approach, we simultaneously fitted Förster radius, linker length, and two DNA geometry parameters (twist and axial rise) after 100 iterations. Here, DNA geometry parameters were allowed to vary slightly to account for unnatural stresses in the nanostructure. As an objective function, the squared sum of the difference between the modeled dye position after triangle construction using given FRET efficiencies (see above) and the expected position given the DNA geometry was used. Supplementary Table 2 denotes ranges, step sizes, and fitted values for all parameters.

#### Extended Figures

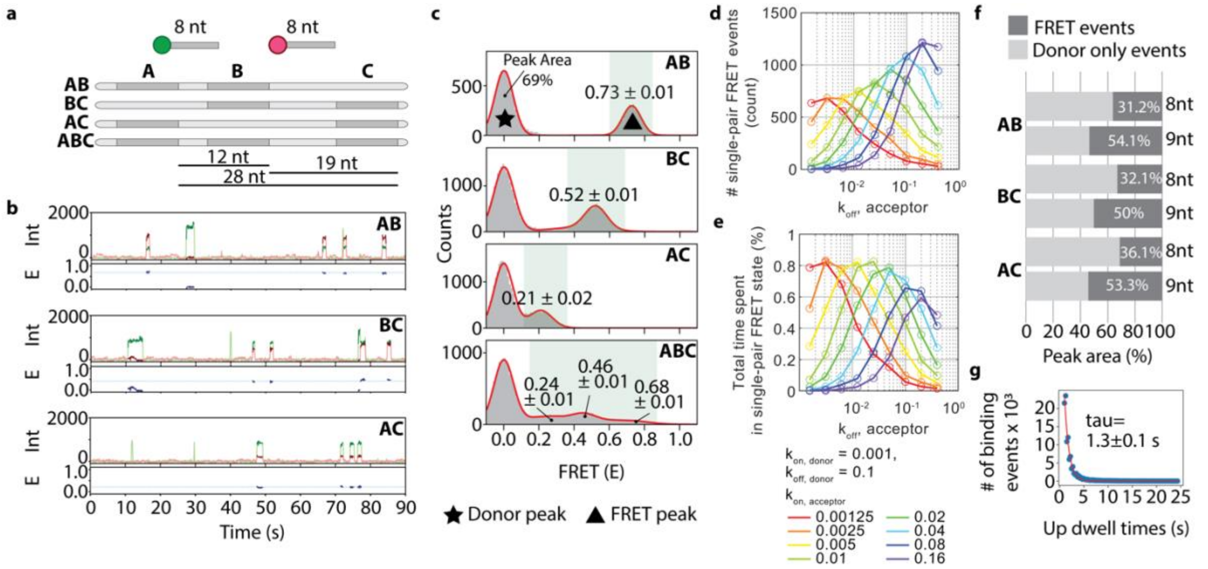


**Extended Fig. 1: Rational design of the linear construct and imager characteristics for iMAX FRET.**

**a**, Schematic representations of the linear DNA constructs. A, B, and C are the positions of identical docking sequences to which 8 nt donor- and acceptor-labeled imagers can bind. The donor to acceptor molar ratio was in a 1:10. The distances between the AB, BC, and AC segments are 12 nt, 19nt, and 28nt, respectively.

**b**, Single-molecule intensity time traces for donor (green), acceptor (red) and FRET (blue) for the linear constructs AB, BC, and AC.

**c**, Single-FRET event histograms from all molecules in a single field of view. Red solid lines are multi-Gaussian fit to the histograms. The three peaks in the ABC construct correspond to the three distances for A-C (0.24 ± 0.01), B-C (0.46 ± 0.01), and A-B (0.68 ± 0.01) (FRET ± SEM). Star designates the donor-only peak whereas the triangle reports the FRET evets peak.

**d-e**, The number of single-pair FRET events (d) and the total time spent (e) of a system with three docking strands, obtained from a series of Monte Carlo simulations with various probe binding kinetic rates. Given the donor binding ($k_{on,\ donor}$ = 0.001) and dissociation ($k_{off,\ donor}$ = 0.1) rates, the number of single-pair FRET events and total time spent within the state changed significantly with the acceptor binding ($k_{on,\ acceptor}$) and dissociation ($k_{off,\ acceptor}$) rates. While the maximum number of events were achieved with higher $k_{on,\ acceptor}$, the maximum time spent started decreasing when $k_{on,\ acceptor}$ was more than 10 times higher than that of the donor. At the optimal 10-times higher $k_{on,\ acceptor}$, $K_{off,\ acceptor}$ should be ~5-10 times lower than that of the donor to maximize the chance of observing single-pair FRET.

**f**, Peak areas for donor only- and single-FRET events for each linear construct are plotted as percentages. Note the increase of 1.5-fold in the FRET events peak area when a longer acceptor imager (9nt) is used instead of an 8nt imager.

**g**, Dwell time histogram of the 9nt acceptor imager binding events (Blue circles). The dwell time determined from single-exponential fit (red line) was 1.3±0.1 s under our experimental conditions (time ± SEM s)

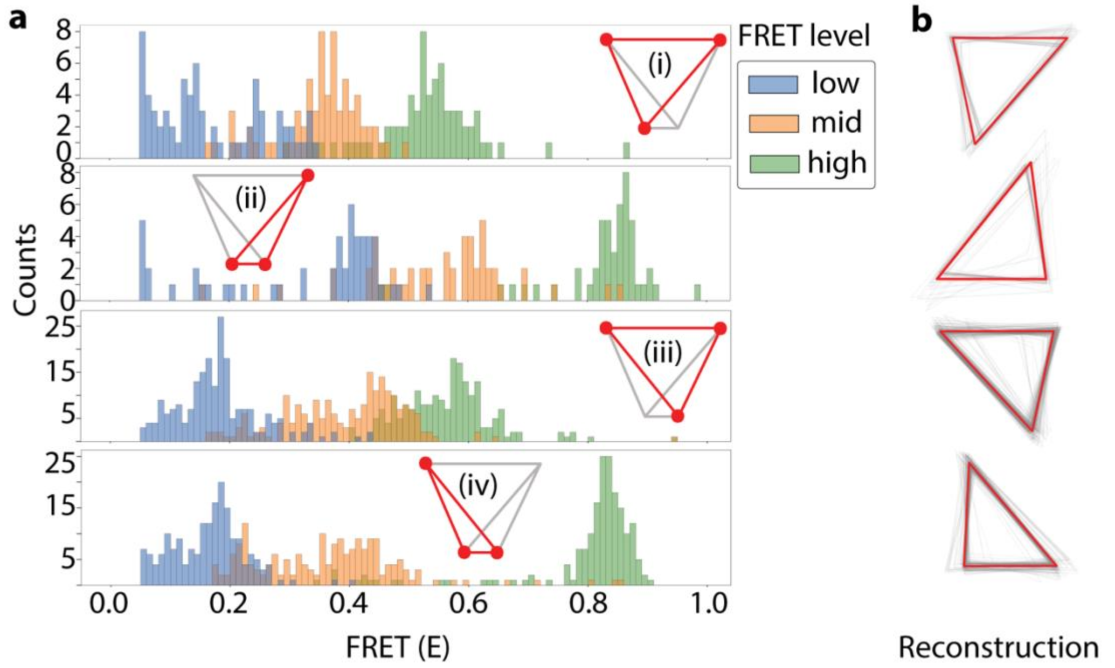


**Extended Fig. 2: Reconstruction of triangles for single-molecule FRET histograms**

a. Histograms displaying per-molecule FRET efficiencies separately for each of the three levels (low, mid, and high) per triangle type in the quadrangular DNA nanostructure (i to iv). Only molecules featuring all three values are shown.

b. Aligned reconstructed triangles for all single molecules (grey) and the average triangle (red) per triangle type.

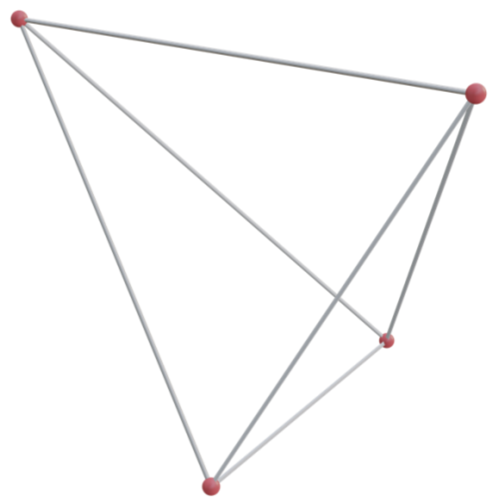

**Extended Fig. 3: 3D reconstruction of the quadrangular DNA nanostructure**
3D reconstruction of the relative dye positions in the nanostructure based on FRET values, revealing its assymetric and staggered nature.

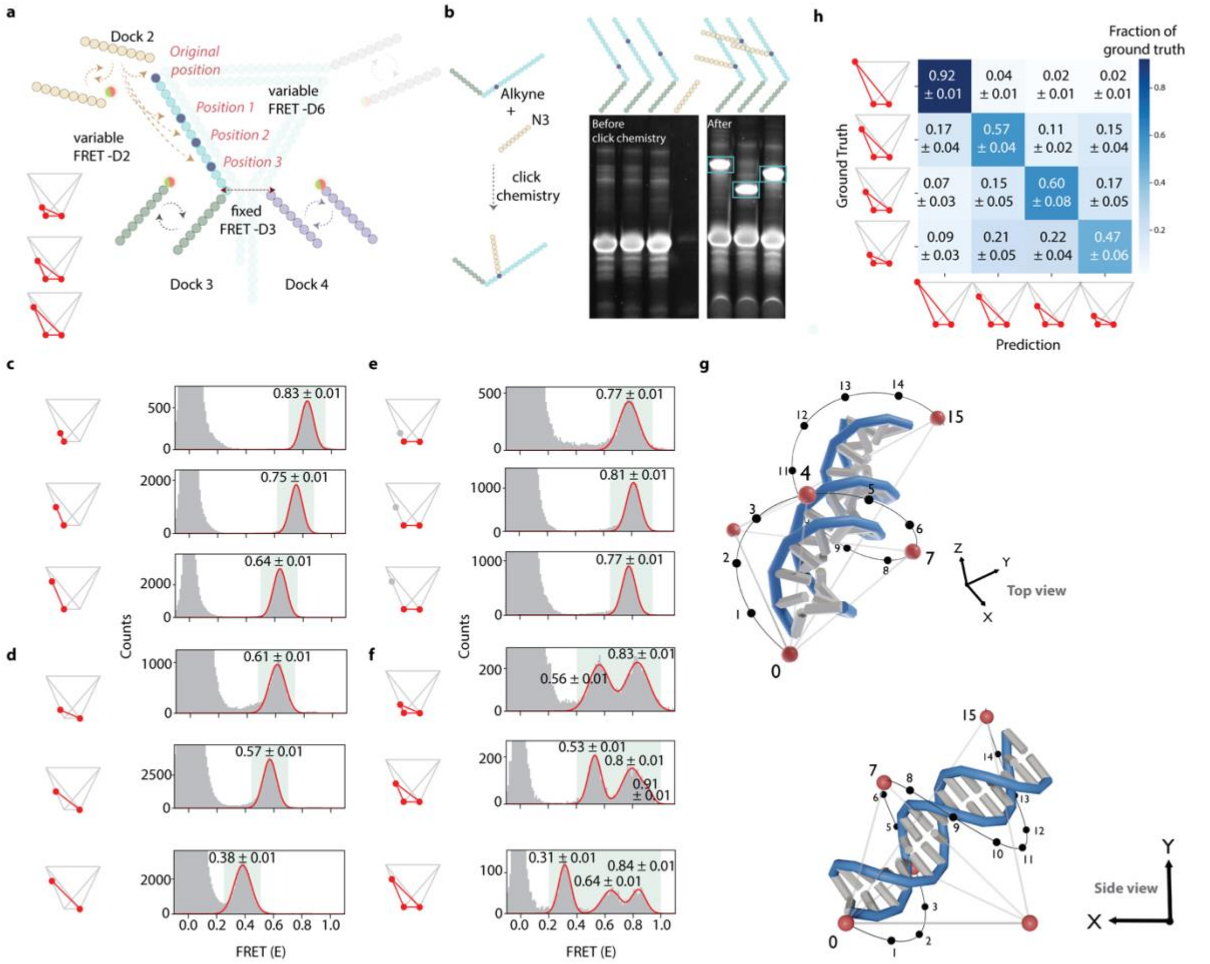


**Extended Fig. 4: iMAX FRET-based analysis of closely related DNA nanostructures**

**a**, In the complex DNA nanostructure, the position of Dock 2 is changed to three different positions giving rise to three FRET-D2 variations.

**b**, Click-chemistry is used to attach an azide-linked Dock 3 to the backbone DNA with an alkyne handle. The clicked DNA products (cyan box) were gel extracted and then the nanostructures were reconstituted by hybridization.

**c**, The FRET changes between the (variable) Dock 2 and fixed Dock 3 are reflected in the change in the differential position change of Dock 2.

**d**, The changes in Dock 2 also changed the distance between sites 2 and 4, as confirmed by the changing FRET values.

**e**, FRET values for the distances, between sites 3 and 4, as expected, remained majorly unaffected.

**f**, The change in FRET values in three points can be similarly recapitulated, for the triangles with imagers and Docks 2,3, and 4. Overall FRET values also shifted for triangles as well for all positions with respect to the original triangle (iv).

**g**, 3D reconstruction of dsDNA strand (blue/white) with dye positions (red spheres) of three triangles with the same base reconstructed from FRET efficiencies. Triangles differed in the position of their third dye, which was located at nucleotides with indices 4, 7, or 15, counted from the base. Förster radius, DNA twist, DNA axial rise, and dye-DNA linker length were optimized using a tree-based Parzen estimator-based approach. Black numbers and dots denote expected dye positions and indices for linkers attached to different nucleotides, based on DNA geometry and linker length. Images rendered at two different view angles were generated in Blender (v3.6).

**h**, our integrated computational approach can differentiate the 3D structures from each other on a single molecule level with up to 60% accuracy.

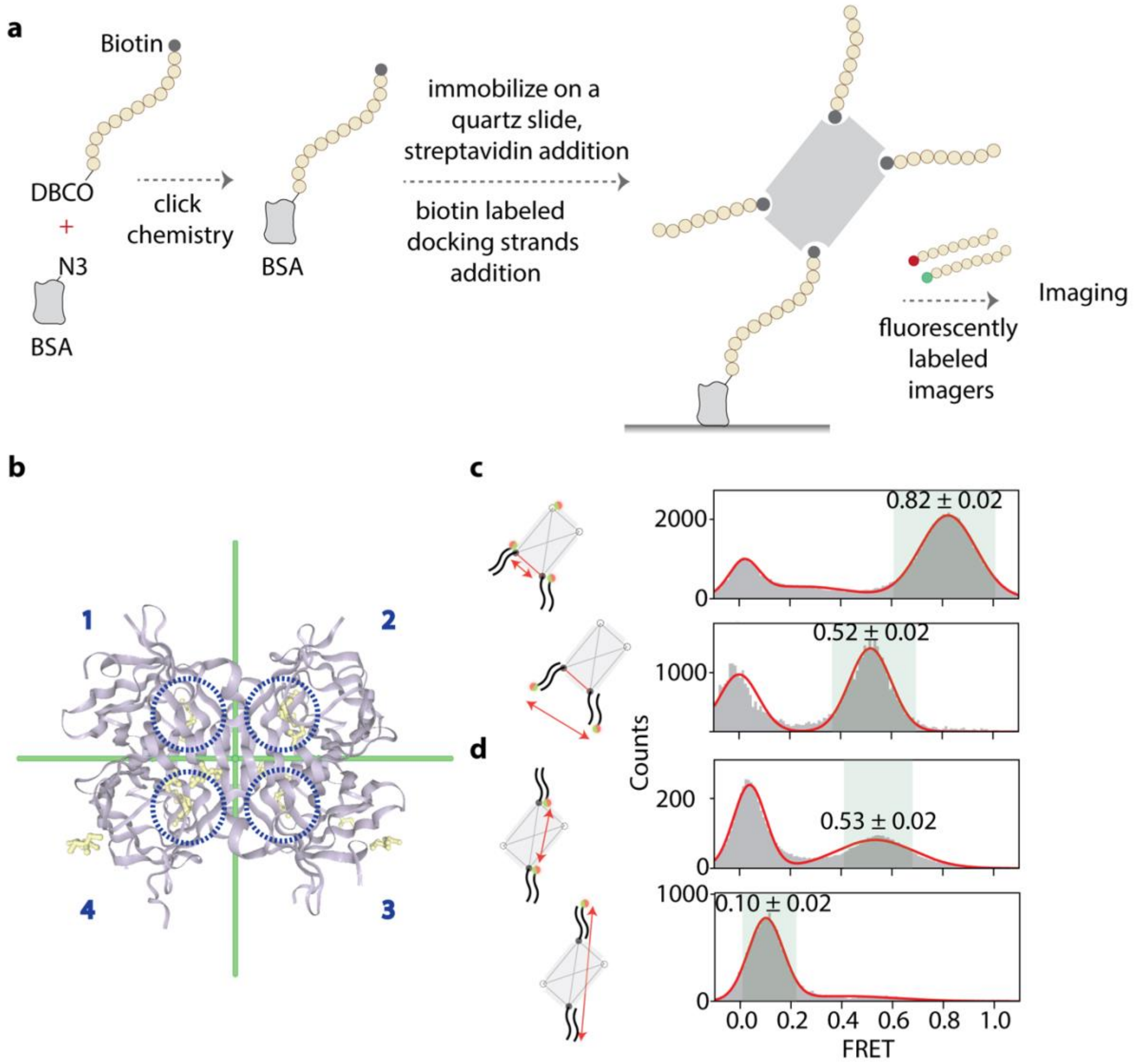


**Extended Fig. 5: Immobilization Scheme of streptavidins and their structural analysis**

**a**, BSA-Azide was immobilized on a quartz slide, conjugated with DNA with a DBCO handle at one end and biotin at the other. The presence of only one Azide per BSA molecule allowed the attachment of one biotin, and thus one streptavidin molecule per BSA molecule. Using this newly developed immobilization scheme, we could ensure that only one pocket is filled with biotin for immobilization and that the remaining 3 pockets are available for binding biotinylated docking sequences for fingerprinting.

**b**, D2 symmetry of the wild-type streptavidin tetramer (from PDB ID: 3RY2). 1,2,3 and 4 designate the numbering of subunits in the tetramer. Biotins (yellow space fills) are highlighted with dashed blue circles.

**c and d**, With the use of imagers for probing as opposed to covalently conjugated dyes generally used in FRET assays, we could modify the location of dye to artificially change the distance between the 2 points. When 3’ instead of 5’ dye-labeled imagers were applied, we could see the relative FRET shift to lower efficiencies corresponding to the new distance, on the divalent structures. 1,2 cis divalent streptavidin shows a change of 0.30 FRET value, while it is 0.43 for the 1,3 trans divalent streptavidin mutants.

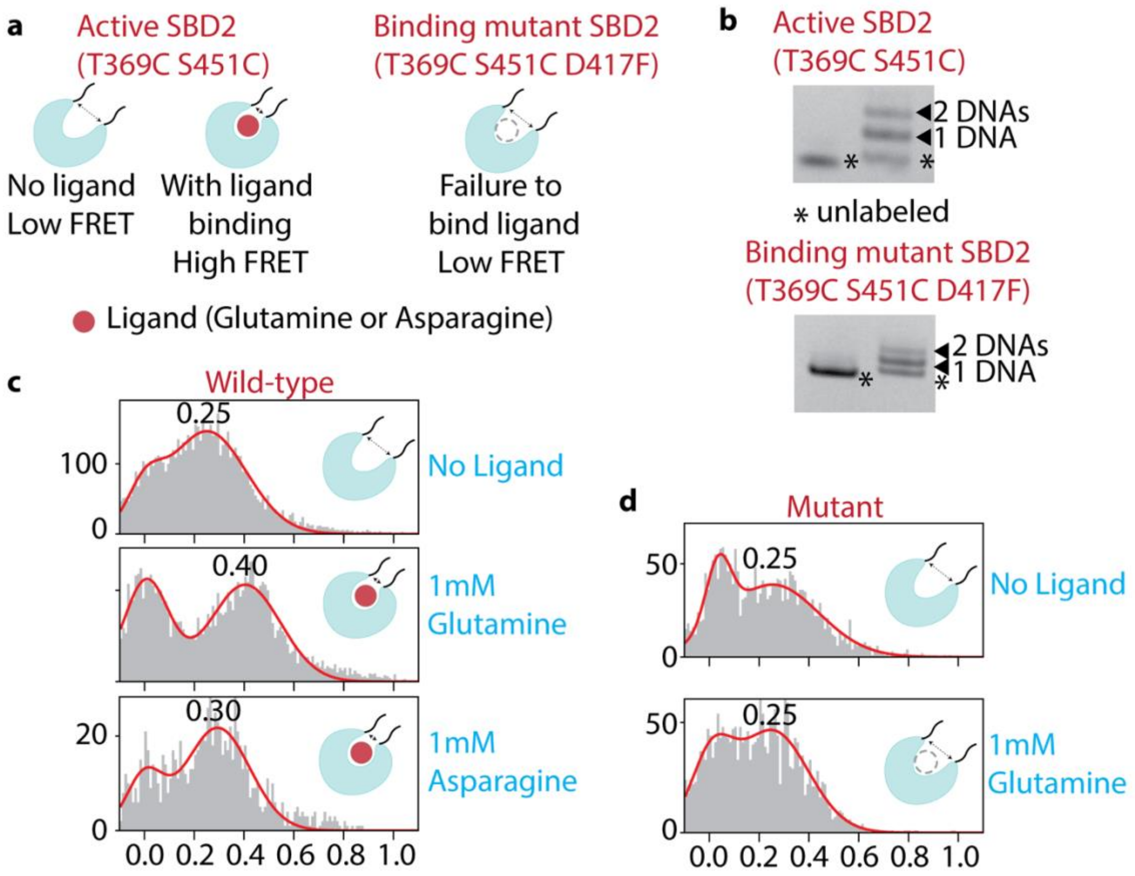


**Extended Fig. 6: Structural analysis of conformational changes in SBD2-ligand complexes**

**a**, 2 mutant SBD2 proteins – active (T369C S451C) and null (T369C S451C D417F). The cysteines are strategically added for DNA labeling. When a cognate ligand is bound, the conformation change results in higher FRET. Whereas, the null mutant retains the low-FRET value due to a lack of ligand binding.

**b**, The SBD2 proteins are labeled with DNA using click chemistry. The ladder pattern suggests the weight shift due to the addition of one or both DNAs attached to the protein.

**c**, The SBD2 protein changes its 0.25 FRET value (no ligand) to 0.40 upon its preferred glutamine ligand binding. When Asparagine is added, it stabilizes at 0.30 FRET.

**d**, The mutant SBD2, due to the inability of ligand binding remains at 0.25 FRET after the application of glutamine.

**Supplementary Tables**

**Supplemetary Table 1: DNA constructs**

| Corresponding Fig. | Description | Sequence (5’-3’) | Modification | Supplier |
|---|---|---|---|---|
| Fig. 2 | Linear construct with POI-A and B docking sequences | ttttttttttttttttttATACATCTAtttATACATCTA | 5’ Biotin | Ella Biotech (GmbH) |
| Fig. 2 | Linear construct with POI-B and C docking sequences | tttttATACATCTAtttttttATACATCTAttttttttt | 5’ Biotin | Ella Biotech (GmbH) |
| Fig. 2 | Linear construct with POI-A and C docking sequences | tttttATACATCTAtttttttttttttttttATACATCTA | 5’ Biotin | Ella Biotech (GmbH) |
| Fig. 2 | Linear construct with POI-A, B and C docking sequences | tttttATACATCTAtttttttATACATCTAtttATACATCTA | 5’ Biotin | Ella Biotech (GmbH) |
| Fig. 2 | Donor imager strand | AGATGTAT | 3’ Cy3 | Ella Biotech (GmbH) |
| Fig. 2 | Acceptor imager strand | AGATGTAT | 3’ Cy5 | Ella Biotech (GmbH) |
| Fig. 2 | Longer acceptor imager strand | TAGATGTAT | 3’ Cy5 | Ella Biotech (GmbH) |
| Fig. 3 | DNA Nanostructure left arm + Dock 1 | AGAGG AGGAT TTCGG TACAC CCGAC AG | - | Ella Biotech (GmbH) |
| Fig. 3 | DNA Nanostructure backbone + Dock 2 | ATTCA TTCTC ATCCT CTGTC GGGTG TACCG TAAGG TGAAT AGTGA CTTTA TACAT CTA | - | Ella Biotech (GmbH) |
| Fig. 3 | DNA Nanostructure biotin strand | CTGAT TGTTA TCGAG GATGA GAATG AATTT TTTTT TTTTT TTT | Biotin – 3’end labeled | Ella Biotech (GmbH) |
| Fig. 3 | DNA Nanostructure right arm + Dock 4 | TCTTC ATTAC TTTTC GATAA CAATC AGGTC ACTAT TCACC TTA | - | Ella Biotech (GmbH) |
| Fig. 3 | DNA Nanostructure Left arm + Dock 2 and Dock 3 | AGAGG AGGAT TTCGG TACAC CCGAC AGTTT TCAAT GTA | - | Ella Biotech (GmbH) |
| Fig. 3 | DNA Nanostructure donor imager strand Dock 1 | AGATGTAT | 3’ Cy3 | Ella Biotech (GmbH) |

| | | | | |
|---|---|---|---|---|
| Fig. 3 | DNA Nanostructure acceptor imager strand Dock 1 | TAGATGTAT | 3’ Cy5 | Ella Biotech (GmbH) |
| Fig. 3 | DNA Nanostructure donor imager strand Dock 2 | TCCTCCT | 5’ Cy3 | Ella Biotech (GmbH) |
| Fig. 3 | DNA Nanostructure acceptor imager strand Dock 2 | TCCTCCTC | 5’ Cy5 | Ella Biotech (GmbH) |
| Fig. 3 | DNA Nanostructure donor imager strand Dock 3 | TACATTGA | 3’ Cy3 | Ella Biotech (GmbH) |
| Fig. 3 | DNA Nanostructure donor imager strand Dock 3 | TACATTGAA | 3’ Cy5 | Ella Biotech (GmbH) |
| Fig. 3 | DNA Nanostructure donor imager strand Dock 4 | AGTAATGA | 5’ Cy3 | Ella Biotech (GmbH) |
| Fig. 3 | DNA Nanostructure acceptor imager strand Dock 4 | AGTAATGAAG | 5’ Cy5 | Ella Biotech (GmbH) |
| Extended Data Fig. 3 | Clickable DNA Nanostructure Left arm + Dock 2 Position 4 | CGGTACACCCGA7AGTTTTCAATGTA | 7= C8-Alkyne-dC | Biomers .net (GmbH) |
| Extended Data Fig. 3 | Clickable DNA Nanostructure Left arm + Dock 2 Position 2 | CGGTA7ACCCGACAGTTTTCAATGTA | 7= C8-Alkyne-dC | Biomers .net (GmbH) |
| Extended Data Fig. 3 | Clickable DNA Nanostructure Left arm + Dock 2 Position 3 | CGGTACACC7GACAGTTTTCAATGTA | 7= C8-Alkyne-dC | Biomers .net (GmbH) |
| Extended Data Fig. 3 | Clickable Dock 2 | AGAGGAGGATTT | 5’ Azide-pro | Biomers .net (GmbH) |
| Fig. 4 | Docking strand for streptavidin WT and mutants | ATACATCTA | 3’ Biotin | Ella Biotech (GmbH) |
| Fig. 4 | Immobilization strand for streptavidin WT and mutants | AAAAGAAAAGAAATACATCTAT | 5’ DBCO, 3’ Biotin | Ella Biotech (GmbH) |
| Extended Data Fig. 4 | Docking strand for proteins – SBD2 WT and mutants | TATACATCTAT | 5’ Azide-pro | Ella Biotech (GmbH) |

**Supplementary Table 2**: **Förster radius fitting parameters**

Ranges, step sizes, and fitted values for all parameters fitted by the tree-based Parzen estimator optimization algorithm. Here, the structure diameter spans the DNA strand diameter and two times the linker length.

| | Min | Max | Step size | Fitted value |
|---|---|---|---|---|
| Förster radius (Å) | 50 | 60 | 0.1 | 53.8 |
| DNA twist (°/bp) | 32 | 40 | 1 | 39 |
| Axial rise (Å/bp) | 2.3 | 5.0 | 0.1 | 4.1 |
| Structure diameter (Å) | 25 | 70 | 0.5 | 38 |
